# Ectomycorrhizas accelerate decomposition to a greater extent than arbuscular mycorrhizas in a northern deciduous forest

**DOI:** 10.1101/2021.02.09.430490

**Authors:** Alexis Carteron, Fabien Cichonski, Etienne Laliberté

## Abstract

It has been proposed that ectomycorrhizal (EcM) fungi slow down decomposition by competing with free-living saprotrophs for organic nutrients and other soil resources (known as the “Gadgil effect”), thereby increasing soil carbon sequestration. As such, this Gadgil effect should depend on soil organic matter age and quality, but this remains unstudied. In addition, the Gadgil effect is not expected to occur in arbuscular mycorrhizal (AM) forests since AM fungi cannot access directly nutrients from soil organic matter, yet few direct comparisons between EcM and AM forests have been made. We performed a two-year reciprocal decomposition experiment of soil organic horizons (litter - L, fragmented - F, humic - H) in adjacent temperate deciduous forests dominated by EcM or AM trees. Litterbags were made of different mesh sizes allowing or excluding ingrowth of external fungal hyphae, which are primarily mycorrhizal in these forests other than for the most-recent superficial litter horizon. As expected, organic matter originating from deeper horizons and from EcM forests was of lower quality (e.g. higher lignin to nitrogen ratios) and decomposed more slowly. However, contrary to the Gadgil effect, organic matter exposed to external fungal hyphae (i.e. primarily mycorrhizal) actually decomposed faster in both forest types, and this effect was strongest in EcM forests, particularly in the F horizon. Unexpectedly, organic matter decomposition was faster in EcM than in AM forests, regardless of organic matter origin. Overall, our study reinforces the view that temperate EcM forests store greater amounts of soil organic carbon than AM forests, but suggests that this is due to factors other than the Gadgil effect.

## Introduction

Forests cover much of the land surface, and represent the largest terrestrial carbon (C) pool globally (Dixon and others 1994; Baldrian 2017). A majority of that C is stored in forest soils, especially in northern forests (Lal 2005; Crowther and others 2019). Soil C storage is controlled by many abiotic and biotic factors such as climate, vegetation, topography and nutrient availability that interact together (Averill and others 2014; Carvalhais and others 2014; Wiesmeier and others 2019). However, belowground biotic factors, such as microorganisms, also play an important role, directly influencing soil C inputs (i.e. litter quantity and quality) and outputs (i.e. decomposition) (Schimel and Schaeffer 2012). For example, soil microorganisms such as fungi can produce recalcitrant organic matter that decomposes slowly or they can produce extracellular enzymes that break down organic matter (Frey 2019). As a result, soil fungi play a major role in forest C cycling (Kubartová and others 2008; Bardgett and Wardle 2010; Orwin and others 2011).

A long-standing hypothesis about the effects of fungi on the soil C cycle is the “Gagdil effect” (Gadgil and Gadgil 1971; Fernandez and Kennedy 2016). This hypothesis suggests ectomycorrhizal (EcM) fungi slow down litter decomposition, potentially due to competition between EcM fungi and free-living saprotrophs for organic nutrients. Because EcM fungi acquire their C in highly labile form via plant hosts (Smith and Read 2008) in exchange for nutrients such as nitrogen and phosphorus, they would leave behind C-rich but nutrient-poor organic matter, potentially favoring soil C accumulation (Read and others 2004; Averill and others 2014). On the other hand, some EcM fungi have the capacity to oxidize organic matter, directly influencing decomposition and indirectly influencing saprotrophic organisms (Lindahl and Tunlid 2015; Verbruggen and others 2017). Saprotrophic fungi could also be impacted by EcM fungi through mycoparatism, antibiosis and alteration of abiotic conditions (Fernandez and Kennedy 2016; Zak and others 2019). The Gagdil effect has only been supported by a few studies but seems to be largely context dependent, for example to litter quality (Smith and Wan 2019) and moisture level (Koide and Wu 2003). Compared to EcM fungi, it is considered that arbuscular mycorrhizal (AM) fungi lack the capacity to produce enzymes that break down organic matter (Tisserant and others 2013; Tedersoo and Bahram 2019). AM fungi would not compete directly with saprotrophic fungi, therefore it is expected that decomposition would be quicker in AM forests compared to EcM forests, but this still remains an open question (Fernandez and Kennedy 2016; Frey 2019). In fact, AM fungi may even enhance directly organic matter decomposition in some cases via a “priming effect”, promoting the activity of free-living saprotrophs (Hodge 2017; Frey 2019). A better understanding of the roles that different mycorrhizal types play in organic matter decomposition is thus needed.

Because fungal types and taxa differ strongly in their vertical distribution, especially in well-stratified soil such as podzols (Dickie and others 2002; Rosling and others 2003; Bahram and others 2015), the strength and direction of the Gadgil effect could vary across soil organic horizons, yet most previous studies have only considered the uppermost litter layer. Strong vertical segregation of fungal guilds occurs across podzol profile: saprotrophic fungi dominate the litter horizon, and can still be abundant in upper organic horizons where mycorrhizal fungi increasingly dominate (Lindahl and others 2007; Clemmensen and others 2015; Carteron and others 2020). It is recognized that overlapping niches between different groups of fungi can generate competition for soil resources (Bödeker and others 2016; Mujic and others 2016). Therefore, the greatest potential for mycorrhizal fungi to inhibit saprotrophs, and thus slow down organic matter decomposition, should lie just below the layer of fresh litter. It has been suggested that these interactions might help to explain differences in the amount and vertical distributions of soil C in EcM systems (Clemmensen and others 2013; Kyaschenko and others 2017) and between EcM- and AM-dominated forests at different depth or horizons (Phillips and others 2013; Soudzilovskaia and others 2015; Craig and others 2018). By competing with saprotrophs for organic nutrients, EcM fungi may promote C accumulation more than AM fungi that cannot directly access these resources. These vertically segregated interactions among fungal guilds need to be better understood because they play an important role in regulating organic matter accumulation (Frey 2019).

Local adaptation to microbial guilds based on soil properties could also be an important factor influencing decomposition via what has been termed the “home-field advantage” (HFA; van der Wal and others 2013) hypothesis. This HFA predicts that litter decomposition is faster in “home soils” due to adaptation of the decomposer community to the chemical composition of the “home litter” (Gholz and others 2000; Austin and others 2014). Using published data on mass loss from 125 reciprocal litter transplants, Veen and others (2015) have shown that this HFA increases decomposition rates by 7.5% on average. However, the strength of the HFA might depend on the context such as plant identity, litter quality and moisture level (Veen and others 2015; Wang and others 2020). Some studies suggest that AM litter shows higher HFA than EcM litter (Midgley and others 2015; Jacobs and others 2018). On the other hand, because EcM litter tends to be more recalcitrant (Keller and Phillips 2019), it could be expected that EcM litter decays faster in EcM forest with saprotrophs better adapted to decompose recalcitrant organic matter. In any case, further investigation is needed to better understand the effect of microbial decomposers driving the HFA depending on litter type and the stage of litter decomposition (Li and others 2020; Lin and others 2020).

The main objective of our study was to assess the impact of EcM and AM strategies on the decomposition of soil organic matter in organic horizons in northern forests. First, we determined stocks of C and nutrients in the upper 20 cm of soil in adjacent forest plots dominated by AM or EcM trees. Then, we performed a litterbag experiment using a reciprocal transplant of organic matter from AM and EcM forest enabling us to isolate site *vs*. organic matter quality effects on decomposition. Litterbags were composed of different mesh size that allowed (44 μm) or excluded (1 μm) ingrowth of fungal hyphae. It is well established that fungal hyphae can move freely across large pore-size (30-50 μm) mesh but prevents in-growth of roots, while small pore-size (0.5-1 μm) mesh further blocks fungal hyphae (Johnson and others 2001; He and others 2004; Teste and others 2009). Since mycorrhizal fungi development requires association with living plant roots, only saprotrophs can develop inside the small-mesh litterbags and access organic matter. Mycorrhizal fungi can colonize organic-rich soil (Lindahl and others 2007; Bunn and others 2019) and, can even be abundant in organic horizons (this system; see Carteron and others 2020). Therefore, large-mesh litterbags allow to follow the decomposition of organic matter in the presence of external fungal hyphae, which are primarily mycorrhizal in these forests other than for the most-recent superficial litter horizon where saprotrophs dominate (Carteron and others 2020). Decomposition of the three upper organic horizons (litter - L, fragmented - F, humic - H) was followed by measuring changes in soil mass, and changes in C and nitrogen (N) over two years. In addition, the fate of C fractions was followed in decomposing L samples and potential access of N by mycorrhiza in the F samples. We hypothesized that the impact of mycorrhizas on organic matter decomposition would differ between AM and EcM forests. More specifically, we expected based on the Gadgil effect hypothesis that EcM forests would store a higher amount of C in the topsoil and show slower organic matter decomposition due to the inhibition of saprotrophs by EcM fungi and lower litter quality, whereas these effects would be smaller in AM forests. In addition, we hypothesized that the slowing down of C cycle by EcM fungi would be strongest in the fragmented (F) horizon where litter-derived organic materials, free-living saprotrophs, mycorrhizal fungi and roots coincide (Clemmensen and others 2013; Cotrufo and others 2015; Carteron and others 2020). Due to microbial adaptations of the decomposer community, we also hypothesized that litter would decompose fastest in their “home” forests relative to “away” forests (Veen and others 2015). Specifically, mass loss of organic matter from AM soil would be highest when incubated in AM forest and mass loss of EcM organic matter highest in EcM forest.

## Material and methods

### Study area and site selection

Our study was conducted in a northern temperate forest at the Université de Montréal’s field station (Station de biologie des Laurentides, Saint-Hippolyte, Québec, Canada). The mean annual temperature is 4.3 °C and total annual precipitation is 1195 mm, with ~25% falling as snow (based on 1981–2010 data, meteorological station #7037310, Saint-Hippolyte). Soils consist of podzols with moder humus formed from Precambrian anorthosite (Bélanger and others 2004; Courchesne and others 2005). We selected ten 20 m × 20 m plots from Carteron and others (2020), either dominated by EcM or AM trees (Table S1), and grouped into five clusters or “blocks” (*n* = 5 blocks, each containing one plot of each of the two mycorrhizal types, EcM and AM). These pairs of EcM-AM sites were clustered together to minimize variation in environmental conditions (e.g. slope, aspect, elevation) within each block. Previous root colonization and molecular analyses on the same sites showed that forests dominated by EcM trees had the highest EcM fungal abundances while forests dominated by AM trees had the highest AM fungal abundances. Carteron and others (2020) also found strong shifts from saprotrophic to mycorrhizal fungal dominance with increasing soil depth in both forest types, especially across surface organic horizons.

### Soil carbon and nutrient stocks

Carbon and nutrient stocks were quantified by measuring C, N, phosphorus (P) concentrations and thickness for all horizons in the upper 20 cm of soil, as reported in Carteron and others (2020). Soil bulk density was measured simultaneously for the five horizons in three randomly-positioned locations replicates per plot using an auger, and values from these locations were averaged across sites. The horizons considered were litter (L), fragmented (F), humic (H), and mineral horizons Ae and B.

### Organic matter collection

In each plot, organic matter samples were collected separately from the three organic horizons, namely: L, F and young H (i.e. most recent layer) from two pits. Samples were homogenized by horizon within each plot. Samples were collected in July 2016. A subsample from each horizon by plot was preserved at 4 °C as inoculum (see below). Another subsample was oven-dried at 60 °C for 72 h and ground for chemical analyses. The rest of the organic matter was air-dried before being used to fill the bags.

### Litterbag design

Litterbags were 15 cm × 15 cm in size and designed to have three compartments (L, F, H; in the same order in which they occur through the soil profile) separated by 44 μm-pore polyethylene mesh (PETEX^®^ 07-40/12; Sefar Inc., Buffalo, NY, USA). Our use of 44 μm-pore mesh ensured that hyphae could grow across compartments within each bag, an important process for decomposition (i.e. to allow for translocation of nutrients and C across horizons), while still keeping L, F, and H horizons separate for later retrieval. The outer mesh of the litterbags was made with either the same 44 μm-pore polyethylene mesh described above or 1 μm-pore mesh from the same material (PETEX^®^ 07-1/2; Sefar Inc., Buffalo, NY, USA). Large pore size (30-50 μm) mesh has been widely used to assess the effect of mycorrhizal hyphal colonization since more than a decade ago (Johnson and others 2001), by excluding fine roots but not fungal hyphae (He and others 2004; Teste 2008). Thus, our litterbags made with 44-μm pore size mesh allow to study decomposition in the presence of mycorrhizal hyphae (and other saprotrophic fungi located outside of the bag). By contrast, the small 1-μm pore size mesh prevents most external fungal hyphae to grow through the litterbag (Teste and others 2006). Because most mycorrhizal hyphae cannot grow within the bag (as mycorrhizal fungi are obligate biotrophs), this bag design allows us to study organic matter decomposition in the absence of mycorrhizal fungi. Litterbags of 50 μm-pore size mesh have been found to allow ingrowth of mycorrhizal fungi (i.e. Teste and others 2006; Sterkenburg and others 2018), which are abundant in our F and H horizons of our plots (Carteron and others 2020). By contrast, free-living saprotrophic fungi should be present in all bags (of 1-μm and 44-μm pore sizes) since all bags were inoculated with horizon-specific organic matter from the same plot prior to being installed in the field. For this reason, while we recognize that the 1-μm mesh bags exclude all external hyphal ingrowth (mycorrhizal and free-living), for simplicity we refer to this treatment as “mycorrhizal exclusion” hereafter since 1-μm mesh bags exclude this particular fungal guild. Mycorrhizal fungal hyphae should be present in the 44-μm mesh bags, being a very important component of the fungal community in the soil other than for the L horizon (Carteron and others 2020). Polyethylene mesh was selected over nylon mesh (e.g. NITEX^®^, Sefar Inc. Buffalo, NY, USA) because it is much more resistant to degradation when buried in soil (Colin and others 1981). Microscope observations showed no evidence that the bags were breached after two years. Previous studies using 0.5 μm-pore size mesh litter bags found that environmental conditions, particularly moisture levels, were similar inside and outside the bags (Allison and others 2013) and, that soil water moved freely across the mesh within minutes (Teste and others 2009). Our own observations confirmed that water moved freely across membranes of both mesh size via capillary action as long as there was contact between the litter inside the bag and the membrane itself; such conditions were maintained throughout the field experiment since the litterbags were buried under the litter layer and secured firmly on the ground. The 1 and 44 μm-pore size mesh have air permeability values of > 95 ±15 l.(m^2^.s)^-1^ at 200 Pa (provided by Sefar inc.). In total, 160 litterbags were used (Fig. S1), from which 90 prevented most hyphal ingrowth of all external fungi. Each bag was stored within a 1-mm mesh nylon bag to provide additional physical protection for the less robust 44- or 1-μm PETEX^®^ mesh.

### Litterbag preparation and collection

Weighed air-dry organic matter was transferred to litterbags (2.85 g for L and 4.75 g for F and H horizons). Horizon specific fresh inoculum (~5 % of dry-weight equivalent) was added to each horizon from the receiving plot to ensure that plot-specific microbial biota, including free-living saprotrophic fungi, could colonize each litterbag. Water content was determined from oven-dried sub-samples at 60 °C for dry-mass inoculum conversion. Filled litterbags were put back *in situ* October 2016, directly on top of the H horizon (with L horizon facing up) and covered by a thin layer of fresh litter. Litterbags were secured on the ground with small stakes and tied together with nylon fishing line to a central stake to facilitate retrieval of bags. Two spatial replicates within each plot were installed. A total of 160 bags were collected after one and two years of residence (i.e. field incubation) for 480 samples analyzed (Fig. S1).

### Soil analysis

Initial subsamples of ground horizons L, F and H were weighed (5.0, 6.0 and 7.0 mg ± 0.2 respectively) and analyzed to estimate C and N contents by dry combustion in a CN analyzer (Vario Micro Cube; Elementar, New-Jersey, United States, dx.doi.org/10.17504/protocols.io.udces2w). The concentrations of soluble cell contents (e.g. non-structural carbohydrates), hemicellulose, cellulose and lignin (% dry weight) were also determined on these initial samples by sequential digestion (Fiber Analyzer 200; ANKOM technology, dx.doi.org/10.17504/protocols.io.yinfude). After one and two years, organic matter samples were retrieved from litterbags, oven-dried at 60 °C for at least 72 h and then weighed to estimate mass loss percentage. These samples were then ground with a cyclone mill (Cyclone Sample Mills, UDY Corporation, Colorado, United States), using a 2-mm screen. Concentrations of C and N were also determined using the method described above. Thirty subsamples of the initial horizons, and all the F horizons after two years of residence were analyzed for δ^15^N with a Micromass model Isoprime 100 isotope ratio mass spectrometer coupled to an Elementar Vario MicroCube elemental analyser in continuous flow mode.

### Statistical analyses

Differences in organic matter stocks among forest types were evaluated using a linear mixed-effects model with forest type (AM or EcM as soil provenance) as a fixed factor and block as a random factor. Horizon was added as fixed factor for the modeling of initial soil chemistry. To predict the changes in mass (within the litterbags), linear mixed-effects models were also used by adding as fixed factors outside fungal hyphae (i.e. size of mesh pore) excluded (1 μm) or not (44 μm). Finally, forest of residence (AM or EcM forest) and time (one or two years) were added as fixed factors to compare decomposition in the two forest types including relevant interactions among fixed factors (see Table S2 for more details). Models were compared using the Akaike information criterion corrected for small sample size (AIC_c_). Validation of the models was done by visual inspection of the residuals. Spatial replicates within one plot were averaged prior to analyses. Eleven bags with damaged mesh were removed from the analysis. Statistical analyses were performed using the R software (R Core Team 2018) and the following packages *dplyr* (Wickham and others 2017), *emmeans* (Lenth 2019), *ggplot2* (Wickham 2016), *ggpubr* (Kassambara 2018), *nlme* (Pinheiro and others 2012). Data and R scripts can be found at https://github.com/alexiscarter/decompo_myco.

## Results

### Organic matter stocks

Stocks of C were higher in EcM forest stands compared to AM stand within the upper 20 cm of soil (one-way analysis of variance, *P* < 0.001; Fig. S2) as observed in the organic horizons (Table 1). Stands dominated by EcM trees stored 14% more C than AM stands in surface soils. The soil C:N ratio also differed among forest types, with higher values in EcM stands (*P* = 0.024; Fig. S3). By contrast, there were no differences in soil C:P ratio among forest types (Fig. S4).

**Table 1.** Initial soil chemistry characteristics: C:N ratio, lignin:N ratio, δ^15^N, thickness and carbon stocks of the upper three horizons (litter - L, fragmented - F, humic - H) of arbuscular mycorrhizal (AM) and ectomycorrhizal (EcM) forest. Means and standard deviation are shown (*n* = 5).

|  | Horizon | AM-dominated | Standard | EcM-dominated | Standard |
| --- | --- | --- | --- | --- | --- |
|  |  | forest | deviation | forest | deviation |
| C:N ratio | L | 22.71 | 2.53 | 27.36 | 1.81 |
|  | F | 20.50 | 1.01 | 22.32 | 1.36 |
|  | H | 18.41 | 0.85 | 20.09 | 1.48 |
| lignin:N ratio | L | 9.68 | 1.40 | 13.05 | 1.74 |
|  | F | 10.70 | 1.69 | 12.57 | 0.99 |
|  | H | 8.62 | 1.30 | 10.50 | 1.50 |
| $\delta^{15}\text{N}$ (‰) | L | -3.26 | 0.36 | -3.16 | 0.63 |
|  | F | -2.48 | 0.40 | -1.96 | 0.48 |
|  | H | -0.72 | 0.59 | -0.74 | 0.36 |
| Thickness (cm) | L | 0.71 | 0.22 | 0.81 | 0.30 |
|  | F | 2.36 | 0.68 | 3.44 | 0.65 |
|  | H | 4.21 | 0.95 | 7.19 | 2.73 |
| Carbon stock | L | 0.89 | 0.28 | 1.05 | 0.43 |
| (kg.m<sup>2</sup>) | F | 2.68 | 0.76 | 4.24 | 0.86 |
|  | H | 2.97 | 0.67 | 6.99 | 4.00 |

### Initial soil chemistry

Soil C, cellulose and hemicellulose concentrations decreased from L to H horizons in both forest types, while lignin was highest in the F horizon and in EcM stands overall (23% in AM forest and 26 % in EcM forest; Table S3). By contrast, soil total [N] increased slightly with soil depth in both forest types (Table S3). As a result, soil C:N and lignin:N ratios were higher in EcM forest for the three organic horizons compared to AM forest (Table S3). δ^15^N values showed similar increases from L to H horizons in both forest types but the F horizon in EcM forest was slightly enriched (but not significantly, *P* = 0.224; Table 1). Horizons tended to be thicker in EcM forest (Table 1).

### Effect of residence on decomposition: AM vs. EcM forests

In AM and EcM stands, older (i.e. deeper) horizons decomposed more slowly than younger ones (Fig. 1). Organic matter loss was slower in the litterbags of 1 μm-pore size mesh in all horizons of both types of forests. However, the slowing down of decomposition due to mycorrhizal fungal exclusion was only statistically significant in the F horizons in stands dominated by AM (−3.7 %, *P* = 0.02; Fig. 1a) and EcM (−4.4 %, *P* = 0.019; Fig. 1b). Differences in the effects of the mycorrhizal exclusion treatments among forest types increased between one and two years of incubation (Fig. 2). Overall, decomposition was slower in AM compared to EcM stands, ranging from −0.8% (*P* > 0.05) of mass loss after one year to −3% (*P* < 0.001) after two years of incubation. After two years, decomposition of organic matter originating from EcM and AM soils was higher in EcM stands (Fig. 2).

**Figure 1.**
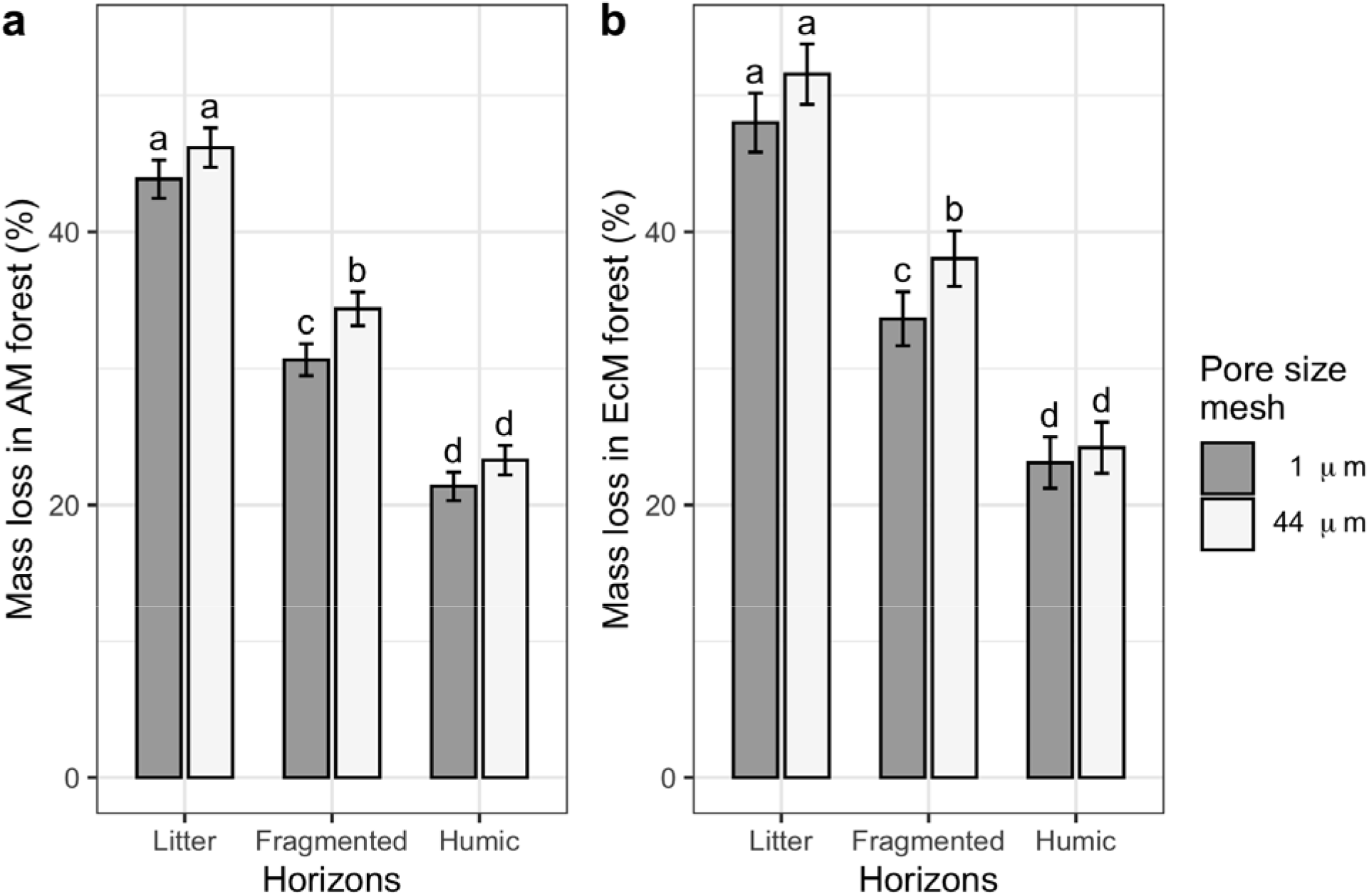
Percentage of mass loss of the three upper horizons incubated for two years in forests dominated by (a) arbuscular mycorrhiza (AM) or (b) ectomycorrhiza (EcM) in litterbags with pore mesh size of 1 μm (grey bars) and 44 μm (white bars). Means ± 1 SE are shown (*n* = 20). Multiple comparison using Tukey′s honestly significant difference post-hoc test, different letters within each panel indicates significant differences (*P*-value < 0.05).

**Figure 2.**
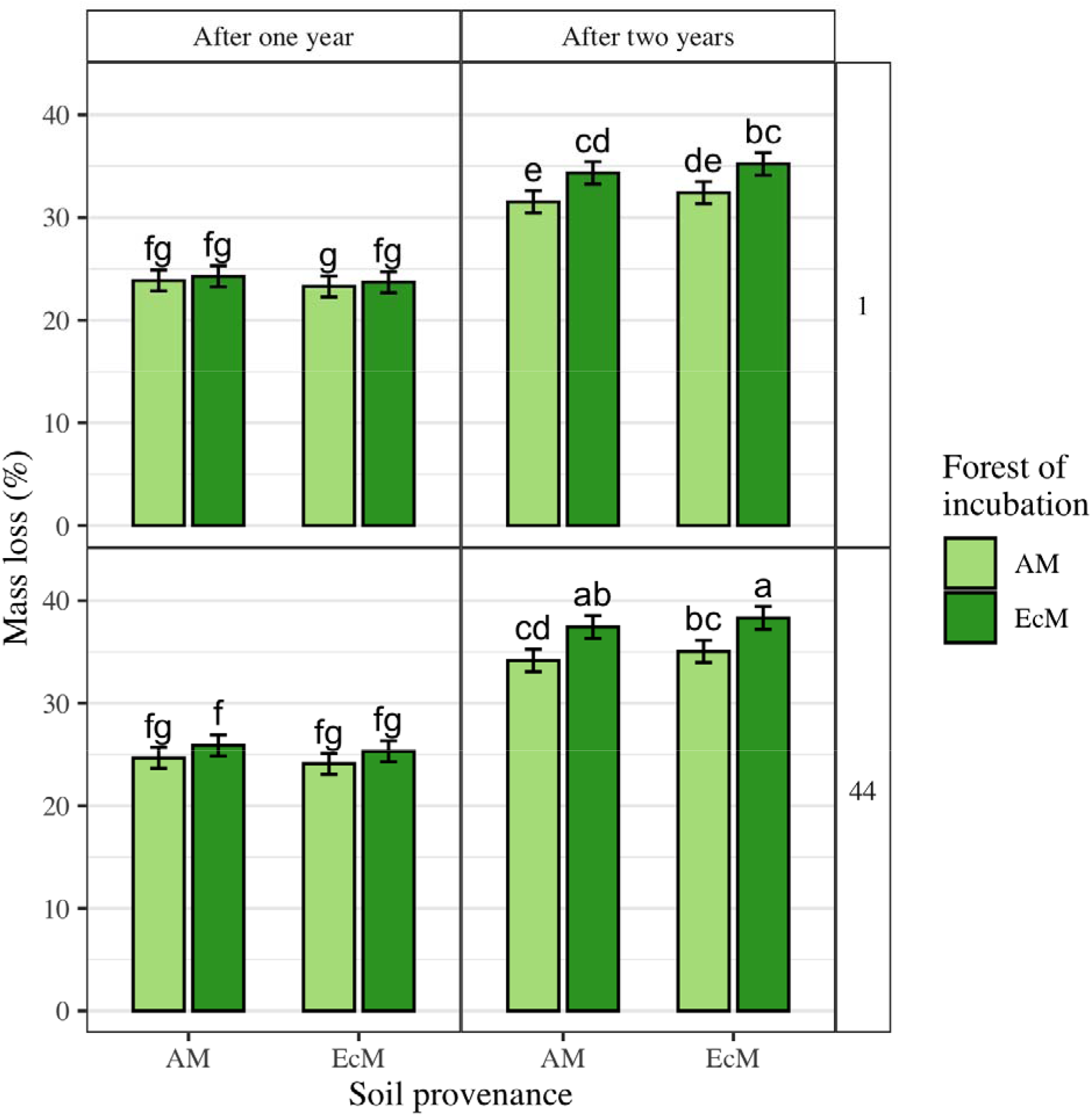
Percentage of mass loss after one and two years of incubation in forests dominated by arbuscular mycorrhiza (AM) or ectomycorrhiza (EcM) in litterbags with pore size mesh of 1 μm (top panels) and 44 μm (bottom panels) and organic matter provenance from AM and EcM. Means ± 1 SE are shown (*n* = 30). Multiple comparison using Tukey′s honestly significant difference post-hoc test, different letters indicates significant differences (*P*-value < 0.05).

### Effect of provenance: AM vs. EcM forests

Litter originating from AM stands decomposed more quickly than EcM litter after one year but this was no longer the case after two years *(P* > 0.05; Fig. S5). While changes in the litter C:N ratio remained similarly low for both soil origins (~1, Fig. S6), the EcM litter showed a clear increase (~1.2) suggesting a lower loss in N compared to C. Similarly, changes in lignin:N ratio were also stronger for EcM litter (Fig. S7).

### Effect of provenance and residence on C fractions and N

Mycorrhizal exclusion did not affect concentrations of soluble contents nor hemicellulose in litter incubated in AM and EcM stands. Compared to EcM stands, decomposition was slower in AM stands for litter cellulose (−4.12 %, *P* = 0.001; Fig. S8) and lignin (−6.56 but *P* > 0.05; Fig. S9). Mycorrhizal exclusion slowed down decomposition of cellulose (−3.79 %, *P* = 0.003) and lignin (−5.13% but *P* > 0.05). Overall, N loss was reduced by mycorrhizal exclusion (−2.83 %, *P* < 0.001) and reduced in AM stands (+2.7 %, *P* < 0.001). After two years, ^15^N enrichment was higher in EcM stands (F horizons only, *P* = 0.046) but effect of mycorrhizal exclusion was rather low (Fig. S10).

## Discussion

No evidence of a Gadgil effect in either forest mycorrhizal type was observed. In fact, the opposite effect was observed, in that decomposition was faster in the presence of EcM or AM fungi than in their absence. Contrary to our hypothesis, decomposition was faster in EcM-than in AM-dominated forests. However, as predicted, decomposition was higher in upper horizons (i.e. “younger” soil), and the net effect of the external fungal network on decomposition was significant in the fragmented (F) horizons. The F horizon is located just below the litter (L), where most decomposition studies tend to focus. Our results suggest that the Gadgil effect is not a universal pattern in EcM forests, and that mycorrhizal fungi may actually accelerate rather than slow down decomposition (Frey 2019). In agreement with these results, we found that decomposition was faster in EcM forests regardless of organic matter origin, suggesting an HFA in EcM but not AM forests.

Several abiotic and biotic factors can impact litter decomposition, such as climate and soil fauna (Hättenschwiler and others 2005; Steidinger and others 2019). However, given the importance of fungi in soil decomposition processes, there has been much interest in exploring the potential effects of interguild fungal interactions over C and nutrient dynamics (Dighton and others 1987; Verbruggen and others 2017). Mycorrhizal fungi can inhibit saprotrophs by competing for nutrients, resulting in slower organic matter decomposition and promotion of C accumulation (Frey 2019). We took advantage of a natural experiment of co-occurring patches of AM and EcM trees under similar environmental conditions but distinct fungal communities and soil chemistry (Carteron and others 2020) to test if contrasting mycorrhizal strategies exerted different control on organic matter decomposition (Phillips and others 2013; Dickie and others 2014). However, contrary to the Gadgil effect hypothesis, our results showed that both EcM and AM fungi accelerate organic matter decomposition in this northern deciduous forest. This might occur if the overall positive effect of mycorrhizal hyphae and other external fungi on decomposition was greater than any potential negative impacts of competition with saprotrophs. In addition, mycorrhizal fungi combined with their local microbial community in EcM forests tended to degrade cellulose and lignin more quickly compared to AM forests. By isolating the effect of mycorrhizas, microbial communities and local environmental conditions, our study shows that decomposition tends to be higher in EcM than AM forests regardless of soil origin and incubation time. Our results challenge the view that EcM fungi slow down litter and soil decomposition compared to AM fungi (Tedersoo and Bahram 2019 and references therein). They also suggest that more attention should be paid to priming *vs*. inhibitory effects of different mycorrhizal types on the decomposition of organic matter (Kuzyakov 2010).

Ectomycorrhizal fungi have traditionally been suggested to slow down litter decomposition via their negative competitive effects on free-living saprotrophs (Gadgil and Gadgil 1971; Fernandez and Kennedy 2016). In our field experiment, we have found that EcM fungi in fact accelerated the decomposition across the three upper organic horizons over two years, particularly in the fragmented (F) horizon. Fernandez & Kennedy (2016) suggested a number of important environmental factors that could modulate the inhibiting impact of EcM fungi on free-living saprotrophs, which might help to explain our results. First, organic matter recalcitrance was relatively low in this broadleaf forest, with lignin:N ratios below 20. Similarly, the C:N ratio was below 30, making N less limiting for saprotrophs compared to other studies where a Gadgil effect was observed (Smith and Wan 2019). Secondly, the studied podzols were well-stratified and exhibited a strong vertical segregation with distinct fungal communities with strong shifts from saprotrophic to mycorrhizal fungal dominance with increasing soil depth (Carteron and others 2020), thereby reducing opportunities for interguild competition. Finally, decreased soil moisture due to EcM fungi can impede decomposition processes (Koide and Wu 2003), but our system is located in a northern temperate forest characterized by a humid continental climate with precipitations throughout the year, where water is not thought to be limiting. At least three other experimental studies have found a positive combined effect of roots and EcM fungi on decomposition (Zhu and Ehrenfeld 1996; Subke and others 2011; Malik 2019). In our case, it is worth noting that the strongest positive net effect was observed in the fragmented horizon where there are: (i) High root colonization by EcM and AM fungi, (ii) high abundance of saprotrophic and mycorrhizal fungi and (iii) high fine root density (Carteron and others 2020). Most studies that have studied the impact of mycorrhizas on decomposition have focus on the most recent litter (L) layer, whereas here we show that important processes occur in deeper (organic) horizons. Our results suggest that vertical stratification should be taken into account to better understand the effect of mycorrhizas on the decomposition process.

Arbuscular mycorrhizal fungi are known to produce compounds that can, for example, alter microbial community or promote soil aggregation thus modulating decomposition rate (e.g. Hodge and others 2001; Gui and others 2017; Xu and others 2018). Decomposition can even be reduced by AM fungi, potentially through antagonistic interactions with free-living saprotrophs (Leifheit and others 2015; Carrillo and others 2016). In our field experiment, we found no evidence of a Gadgil effect exerted by AM fungi that would counterbalance their positive impacts on decomposition. As expected, decomposition in the upper three organic horizons in AM forest was not reduced with the 1-μm mesh bags (in which mycorrhizal hyphae were excluded), but in fact tended to increase. Given that AM fungi lack a strong degradation machinery (Tedersoo and Bahram 2019), our results support the view that priming of organic matter decomposition might be an important nutrient acquisition strategy for them (Wurzburger and Brookshire 2017). Greater priming in AM systems may result from AM fungal necromass and the lack of genetic capacity from AM fungi to directly access organic nutrients (Frey 2019). It is worth noting that with the 1-μm mesh bags, decomposition in AM forests tended to be slower than in EcM forests, suggesting that their free-living saprotrophic communities have different capacities to degrade organic matter (see results after two years in Fig. 2). Microbial communities in AM forest might be less efficient at degrading organic matter due to their more easily-decomposed litter contrary to what has been observed for other AM systems in microcosm experiments (Taylor and others 2016). Evaluating the response of the saprotrophic community using molecular tools over long-term experiment would be an interesting way to better understand decomposition processes *in situ*, in order to complement studies that focus on laboratory manipulations of mycorrhizal abundance (Verbruggen and others 2017). It would also allow us to experimentally assess if the abundance of saprotrophs shifts in deeper horizons when AM and EcM fungi are excluded (e.g. Lindahl and others 2010; Sietiö and others 2019).

Leaf litter decomposition rates are known to be positively linked with initial N concentration and inversely with lignin (Prescott 2005; Berg and McClaugherty 2014). Overall net effect of mycorrhizas over decomposition is known to be controlled by substrate quality and local microbial community composition (Fernandez and others 2019; Smith and Wan 2019). As expected, we found a strong effect of soil depth with deeper (i.e. “older”) horizons decomposing more slowly. In general, the litter of EcM-associated trees tend to have a lower quality than AM trees such as lower C:N and lignin:N ratios (Lin and others 2017), which may drive soil C accumulation in the short-term. In our field sites, litter in EcM stands had lower quality compared to AM stands. The EcM stands were mostly composed of American beech. However, American beech litter is less recalcitrant than many conifers (Moore and others 1999), which may explain discrepancies with other studies from coniferous EcM forests in which Gadgil effects have been observed (Fernandez and Kennedy 2016; Smith and Wan 2019). In temperate forests, AM plants tend to produce leaf litter that decomposes more rapidly *in situ* than that of EcM plants (Keller and Phillips 2019). Similarly, litter originating from AM patches decomposed more quickly than EcM litter after one year but interestingly, this was not the case after two years in our experiment. This is consistent with previous studies showing that sugar maple leaf litter tend to decomposes more quickly during the first years after senescence (McHale and others 1998; Lovett and others 2016), but tends to be more similar to American beech after several years (within standard error range, see Lovett and others (2016). Community of decomposers in EcM forests may be efficient at decomposing recalcitrant organic matter (Fernandez and others 2019). Contrary to the results of Midgley and others (2015) obtained from another study system, we observed HFA in EcM forests but not in AM forests. The efficiency of the microbial decomposers present in EcM soil for decomposing organic matter may be high regardless of litter type and quality. Furthermore, we found no evidence that fragmented (F) and humic (H) horizons in AM stands decomposed faster than the same horizons in EcM stands (but see Jacobs and others 2018). Taken together, these results suggest that the significant impact of initial litter chemistry on decomposition diminishes after the first year of decomposition and that microbial decomposer community may adapt to “home” substrate quality.

Reducing decomposer diversity reduces litter decomposition rate (Handa and others 2014; Li and others 2020), but this effect is context-dependent and the effect of soil fauna is variable across focal species (Makkonen and others 2012). Smaller mesh size are known to reduce the potential diversity of soil fauna that are important for decomposition processes (Hättenschwiler and others 2005). In our study, patches were mainly composed of American beech or sugar maple, and previous studies indicate that maple litter is generally preferred over beech litter by the soil fauna (Hättenschwiler and Bretscher 2001; Jacob and others 2010). However, the difference in decomposition between American beech and sugar maple seems to decrease over time (Lovett and others 2016) and to be dependent on stand type of incubation (Côté and Fyles 1994). Unlike most studies, we used litterbags that were designed to follow decomposition of the upper three organic horizons while avoiding soil trenching and tree girdling which confound the effects of roots and mycorrhizal fungi (Fernandez and Kennedy 2016). Trenching is the historical and most widely used method to test the Gadgil effect, but it is known to directly affect soil drainage, increase soil moisture by impeding root water uptake and strongly disturb the system (Gadgil and Gadgil 1971; Fisher and Gosz 1986; Fernandez and Kennedy 2016). Tree girdling is the most extreme alternative as it kills trees, also preventing further research on the same site. In our experiment, the initial disturbance may have increased labile C but the persistence of this effect after two years was assumed to be rather limited. Furthermore, the observed effects of our exclusion treatment on mass loss increased between the first and second years of incubation suggesting persistent biological effects. Decreasing mesh size might have decreased soil moisture but we observed no impact on litter soluble content losses suggesting a rather low effect caused by mesh size, at least on the most labile fractions of C. It is possible that the use of litterbags with small mesh size limited the exposure to biophysical perturbations, which might hamper mass loss (Prescott 2005; Berg and McClaugherty 2014), but this was common to all treatments. To better predict soil C processes and stocks, more research may be needed to understand how interaction between mycorrhizas, soil fauna, plant inputs and variables such as soil moisture, or bulk density impact decomposition (Lin and others 2017).

Our sampling design allowed us to spatially distinguish decomposition processes in the upper three horizons and assess the fate of young to older organic matter overcoming some limits of short-term experiments. The overall net effect of mycorrhizas on decomposition was positive regardless of mycorrhizal type, but varied throughout the soil profile. Further analyses would allow us to better understand if the greater decomposition could be due to higher microbial biomass inside the 44 μm-pore mesh bags, leading to higher enzymatic activities. As expected from previous studies (e.g., Averill and others 2014; Soudzilovskaia and others 2019), C stocks were greater in EcM stands compared to neighboring AM stands in this northern temperate forest even though decomposition was greater in EcM soils, and positively influenced by the broader fungal network. This indicates the potential importance of others factors such as litter quantity, soil fauna and moisture level in regulating C dynamics. The quality and composition of litter is important for short-term C release, but the microbial community, including root-associated fungi and mycorrhizal-associated organisms (Netherway and others 2020), potentially have a strong impact on a longer-term which is important for C sequestration (Cotrufo and others 2015). Overall, our study shows that forests dominated by different mycorrhizal strategies have distinct impacts in soil organic matter dynamics. The EcM forests store higher soil organic C but support microbial decomposer communities that are more efficient at degrading organic matter than those of adjacent AM forests, rejecting the Gadgil effect as a driver of C accumulation in the northern temperate forests.

## Acknowledgements

We would like to thank the staff from the Station de biologie des Laurentides (SBL) of Université de Montréal for facilitating the field work. Funding, including scholarships to AC, was provided by Discovery Grants to EL (RGPIN-2014-06106, RGPIN-2019-04537) by the Natural Sciences and Engineering Research Council of Canada (NSERC) as well as a “Nouveau Chercheur” grant (2016-NC-188823) by the Fonds de recherche du Québec sur la Nature et technologies (FRQNT). AC would like to sincerely thank the following institutions for providing generous scholarships: FRQNT (Dossier 272522), Institut de recherche en biologie végétale, Centre de la science de la biodiversité du Québec, Centre d′étude de la forêt and Université de Montréal through the “Bourse d′excellence Hydro-Québec”.

## Authors′ contributions

EL and AC conceived the ideas and designed methodology; AC and FC collected and analyzed the data; AC and EL interpreted the results; AC led the writing of the manuscript. All authors contributed critically to the drafts and gave final approval for publication.

## Conflict of Interest

The authors declare that they have no conflict of interest.

## Notes

### Competing Interest Statement

The authors have declared no competing interest.

## Literature Cited

Allison SD, Lu Y, Weihe C, Goulden ML, Martiny AC, Treseder KK, Martiny JBH. 2013. Microbial abundance and composition influence litter decomposition response to environmental change. Ecology 94:714–25.

Austin AT, Vivanco L, González-Arzac A, Pérez LI. 2014. There’s no place like home? An exploration of the mechanisms behind plant litter–decomposer affinity in terrestrial ecosystems. New Phytologist 204:307–14.

Averill C, Turner BL, Finzi AC. 2014. Mycorrhiza-mediated competition between plants and decomposers drives soil carbon storage. Nature 505:543–5.

Bahram M, Peay KG, Tedersoo L. 2015. Local-scale biogeography and spatiotemporal variability in communities of mycorrhizal fungi. New Phytol 205:1454–63.

Baldrian P. 2017. Forest microbiome: diversity, complexity and dynamics. FEMS Microbiol Rev 41:109–30.

Bardgett RD, Wardle DA. 2010. Aboveground-belowground linkages: Biotic interactions, ecosystem processes, and global change. OUP Oxford

Bélanger N, Côté B, Fyles JW, Courchesne F, Hendershot WH. 2004. Forest regrowth as the controlling factor of soil nutrient availability 75 years after fire in a deciduous forest of Southern Quebec. Plant and Soil 262:363–272.

Berg B, McClaugherty C. 2014. Plant Litter: Decomposition, Humus Formation, Carbon Sequestration. 3rd ed. Berlin Heidelberg: Springer-Verlag.

Bödeker ITM, Lindahl BD, Olson Å, Clemmensen KE. 2016. Mycorrhizal and saprotrophic fungal guilds compete for the same organic substrates but affect decomposition differently. Funct Ecol 30:1967–78.

Bunn RA, Simpson DT, Bullington LS, Lekberg Y, Janos DP. 2019. Revisiting the ‘direct mineral cycling’ hypothesis: arbuscular mycorrhizal fungi colonize leaf litter, but why? The ISME Journal 13:1891.

Carrillo Y, Dijkstra FA, LeCain D, Pendall E. 2016. Mediation of soil C decomposition by arbuscular mycorrizhal fungi in grass rhizospheres under elevated CO_2_. Biogeochemistry 127:45–55.

Carteron A, Beigas M, Joly S, Turner BL, Laliberté E. 2020. Temperate forests dominated by arbuscular or ectomycorrhizal fungi are characterized by strong shifts from saprotrophic to mycorrhizal fungi with increasing soil depth. Microb Ecol:1–14.

Carvalhais N, Forkel M, Khomik M, Bellarby J, Jung M, Migliavacca M, Mu M, Saatchi S, Santoro M, Thurner M, Weber U, Ahrens B, Beer C, Cescatti A, Randerson JT, Reichstein M. 2014. Global covariation of carbon turnover times with climate in terrestrial ecosystems. Nature 514:213–7.

Clemmensen KE, Bahr A, Ovaskainen O, Dahlberg A, Ekblad A, Wallander H, Stenlid J, Finlay RD, Wardle DA, Lindahl BD. 2013. Roots and associated fungi drive long-term carbon sequestration in boreal forest. Science 339:1615–8.

Clemmensen KE, Finlay RD, Dahlberg A, Stenlid J, Wardle DA, Lindahl BD. 2015. Carbon sequestration is related to mycorrhizal fungal community shifts during long-term succession in boreal forests. New Phytol 205:1525–36.

Colin G, Cooney JD, Carlsson DJ, Wiles DM. 1981. Deterioration of plastic films under soil burial conditions. J Appl Polym Sci 26:509–19.

Côté B, Fyles JW. 1994. Leaf litter disappearance of hardwood species of southern Québec: Interaction between litter quality and stand type. Écoscience 1:322–8.

Cotrufo MF, Soong JL, Horton AJ, Campbell EE, Haddix ML, Wall DH, Parton WJ. 2015. Formation of soil organic matter via biochemical and physical pathways of litter mass loss. Nature Geoscience 8:776–9.

Courchesne F, Côté B, Fyles JW, Hendershot WH, Biron PM, Roy AG, Turmel M-C. 2005. Recent changes in soil chemistry in a forested ecosystem of southern Québec, Canada. Soil Science Society of America Journal 69:1298.

Craig ME, Turner BL, Liang C, Clay K, Johnson DJ, Phillips RP. 2018. Tree mycorrhizal type predicts within-site variability in the storage and distribution of soil organic matter. Global Change Biology 24:3317–30.

Crowther TW, Hoogen J van den, Wan J, Mayes MA, Keiser AD, Mo L, Averill C, Maynard DS. 2019. The global soil community and its influence on biogeochemistry. Science 365:eaav0550.

Dickie IA, Koele N, Blum JD, Gleason JD, McGlone MS. 2014. Mycorrhizas in changing ecosystems. Botany 92:149–60.

Dickie IA, Xu B, Koide RT. 2002. Vertical niche differentiation of ectomycorrhizal hyphae in soil as shown by T-RFLP analysis. New Phytologist 156:527–35.

Dighton J, Thomas ED, Latter PM. 1987. Interactions between tree roots, mycorrhizas, a saprotrophic fungus and the decomposition of organic substrates in a microcosm. Biol Fert Soils 4:145–50.

Dixon RK, Solomon AM, Brown S, Houghton RA, Trexier MC, Wisniewski J. 1994. Carbon pools and flux of global forest ecosystems. Science 263:185–90.

Fernandez CW, Kennedy PG. 2016. Revisiting the ‘Gadgil effect’: do interguild fungal interactions control carbon cycling in forest soils? New Phytol 209:1382–94.

Fernandez CW, See CR, Kennedy PG. 2019. Decelerated carbon cycling by ectomycorrhizal fungi is controlled by substrate quality and community composition. New Phytologist 226:569–82.

Fisher FM, Gosz JR. 1986. Effects of trenching on soil processes and properties in a New Mexico mixed-conifer forest. Biol Fert Soils 2:35–42.

Frey SD. 2019. Mycorrhizal fungi as mediators of soil organic matter dynamics. Annu Rev Ecol Evol Syst 50:237–59.

Gadgil RL, Gadgil PD. 1971. Mycorrhiza and litter decomposition. Nature 233:133–133.

Gholz HL, Wedin DA, Smitherman SM, Harmon ME, Parton WJ. 2000. Long-term dynamics of pine and hardwood litter in contrasting environments: toward a global model of decomposition. Global Change Biology 6:751–65.

Gui H, Hyde K, Xu J, Mortimer P. 2017. Arbuscular mycorrhiza enhance the rate of litter decomposition while inhibiting soil microbial community development. Scientific Reports 7:42184.

Handa IT, Aerts R, Berendse F, Berg MP, Bruder A, Butenschoen O, Chauvet E, Gessner MO, Jabiol J, Makkonen M, McKie BG, Malmqvist B, Peeters ETHM, Scheu S, Schmid B, Ruijven J van, Vos VCA, Hättenschwiler S. 2014. Consequences of biodiversity loss for litter decomposition across biomes. Nature 509:218–21.

Hättenschwiler S, Bretscher D. 2001. Isopod effects on decomposition of litter produced under elevated CO_2_, N deposition and different soil types. Global Change Biology 7:565–79.

Hättenschwiler S, Tiunov AV, Scheu S. 2005. Biodiversity and litter decomposition in terrestrial ecosystems. Annual Review of Ecology, Evolution, and Systematics 36:191–218.

He X, Critchley C, Ng H, Bledsoe C. 2004. Reciprocal N (15NH4+ or 15NO3-) transfer between nonN2-fixing Eucalyptus maculata and N2-fixing Casuarina cunninghamiana linked by the ectomycorrhizal fungus Pisolithus sp. New Phytologist 163:629–40.

Hodge A. 2017. Chapter 8 - Accessibility of inorganic and organic nutrients for mycorrhizas. In: Mycorrhizal Mediation of Soil. Elsevier. pp 129–48.https://www.sciencedirect.com/science/article/pii/B9780128043127000085

Hodge A, Campbell CD, Fitter AH. 2001. An arbuscular mycorrhizal fungus accelerates decomposition and acquires nitrogen directly from organic material. Nature 413:297–9.

Jacob M, Viedenz K, Polle A, Thomas FM. 2010. Leaf litter decomposition in temperate deciduous forest stands with a decreasing fraction of beech (*Fagus sylvatica*). Oecologia 164:1083–94.

Jacobs LM, Sulman BN, Brzostek ER, Feighery JJ, Phillips RP. 2018. Interactions among decaying leaf litter, root litter and soil organic matter vary with mycorrhizal type. Journal of Ecology 106:502–13.

Johnson D, Leake JR, Read DJ. 2001. Novel in-growth core system enables functional studies of grassland mycorrhizal mycelial networks. New Phytologist 152:555–62.

Kassambara A. 2018. ggpubr: ‘ggplot2’ Based Publication Ready Plots. https://CRAN.R-project.org/package=ggpubr https://CRAN.R-project.org/package=ggpubr

Keller AB, Phillips RP. 2019. Leaf litter decay rates differ between mycorrhizal groups in temperate, but not tropical, forests. New Phytologist 222:556–64.

Koide RT, Wu T. 2003. Ectomycorrhizas and retarded decomposition in a *Pinus resinosa* plantation. New Phytologist 158:401–7.

Kubartová A, Ranger J, Berthelin J, Beguiristain T. 2008. Diversity and decomposing ability of saprophytic fungi from temperate forest litter. Microb Ecol 58:98–107.

Kuzyakov Y. 2010. Priming effects: Interactions between living and dead organic matter. Soil Biology and Biochemistry 42:1363–71.

Kyaschenko J, Clemmensen KE, Karltun E, Lindahl BD. 2017. Below-ground organic matter accumulation along a boreal forest fertility gradient relates to guild interaction within fungal communities. Ecol Lett 20:1546–55.

Lal R. 2005. Forest soils and carbon sequestration. Forest Ecology and Management 220:242–58.

Leifheit EF, Verbruggen E, Rillig MC. 2015. Arbuscular mycorrhizal fungi reduce decomposition of woody plant litter while increasing soil aggregation. Soil Biology and Biochemistry 81:323–8.

Lenth R. 2019. emmeans: Estimated Marginal Means, aka Least-Squares Means. https://CRAN.R-project.org/package=emmeans https://CRAN.R-project.org/package=emmeans

Li Y, Veen GF (Ciska), Hol WHG, Vandenbrande S, Hannula SE, ten Hooven FC, Li Q, Liang W, Bezemer TM. 2020. ‘Home’ and ‘away’ litter decomposition depends on the size fractions of the soil biotic community. Soil Biology and Biochemistry 144:107783.

Lin D, Dou P, Yang G, Qian S, Wang H, Zhao L, Yang Y, Mi X, Ma K, Fanin N. 2020. Home-field advantage of litter decomposition differs between leaves and fine roots. New Phytologist 227:995–1000.

Lin G, McCormack ML, Ma C, Guo D. 2017. Similar below-ground carbon cycling dynamics but contrasting modes of nitrogen cycling between arbuscular mycorrhizal and ectomycorrhizal forests. New Phytologist 213:1440–51.

Lindahl BD, de Boer W, Finlay RD. 2010. Disruption of root carbon transport into forest humus stimulates fungal opportunists at the expense of mycorrhizal fungi. The ISME Journal 4:872–81.

Lindahl BD, Ihrmark K, Boberg J, Trumbore SE, Högberg P, Stenlid J, Finlay RD. 2007. Spatial separation of litter decomposition and mycorrhizal nitrogen uptake in a boreal forest. New Phytologist 173:611–20.

Lindahl BD, Tunlid A. 2015. Ectomycorrhizal fungi – potential organic matter decomposers, yet not saprotrophs. New Phytol 205:1443–7.

Lovett GM, Arthur MA, Crowley KF. 2016. Effects of calcium on the rate and extent of litter decomposition in a northern hardwood forest. Ecosystems 19:87–97.

Makkonen M, Berg MP, Handa IT, Hättenschwiler S, Ruijven J van, Bodegom PM van, Aerts R. 2012. Highly consistent effects of plant litter identity and functional traits on decomposition across a latitudinal gradient. Ecology Letters 15:1033–41.

Malik RJ. 2019. No “Gadgil effect”: Temperate tree roots and soil lithology are effective predictors of wood decomposition. Forest Pathology 49:e12506.

McHale PJ, Mitchell MJ, Bowles FP. 1998. Soil warming in a northern hardwood forest: trace gas fluxes and leaf litter decomposition. Can J For Res 28:1365–72.

Midgley MG, Brzostek E, Phillips RP. 2015. Decay rates of leaf litters from arbuscular mycorrhizal trees are more sensitive to soil effects than litters from ectomycorrhizal trees. Journal of Ecology 103:1454–63.

Moore TR, Trofymow JA, Taylor B, Prescott C, Camiré C, Duschene L, Fyles J, Kozak L, Kranabetter M, Morrison I, Siltanen M, Smith S, Titus B, Visser S, Wein R, Zoltai S. 1999. Litter decomposition rates in Canadian forests. Global Change Biology 5:75–82.

Mujic AB, Durall DM, Spatafora JW, Kennedy PG. 2016. Competitive avoidance not edaphic specialization drives vertical niche partitioning among sister species of ectomycorrhizal fungi. New Phytologist 209:1174–83.

Netherway T, Bengtsson J, Krab EJ, Bahram M. 2020. Biotic interactions with mycorrhizal systems as extended nutrient acquisition strategies shaping forest soil communities and functions. Basic and Applied Ecology: 10.1016/j.baae.2020.10.002.

Orwin KH, Kirschbaum MUF, St John MG, Dickie IA. 2011. Organic nutrient uptake by mycorrhizal fungi enhances ecosystem carbon storage: a model-based assessment. Ecology Letters 14:493–502.

Phillips RP, Brzostek E, Midgley MG. 2013. The mycorrhizal-associated nutrient economy: a new framework for predicting carbon–nutrient couplings in temperate forests. New Phytol 199:41–51.

Pinheiro J, Bates D, DebRoy S, Sarkar D, Team RC. 2012. nlme: Linear and nonlinear mixed effects models. R package version 3.

Prescott CE. 2005. Do rates of litter decomposition tell us anything we really need to know? Forest Ecology and Management 220:66–74.

R Core Team. 2018. R: A Language and Environment for Statistical Computing. Vienna, Austria: R Foundation for Statistical Computing https://www.R-project.org/

Read DJ, Leake JR, Perez-Moreno J. 2004. Mycorrhizal fungi as drivers of ecosystem processes in heathland and boreal forest biomes. Can J Bot 82:1243–63.

Rosling A, Landeweert R, Lindahl BD, Larsson K-H, Kuyper TW, Taylor AFS, Finlay RD. 2003. Vertical distribution of ectomycorrhizal fungal taxa in a podzol soil profile. New Phytologist 159:775–83.

Schimel JP, Schaeffer SM. 2012. Microbial control over carbon cycling in soil. Front Microbiol 3:348.

Sietiö O-M, Santalahti M, Putkinen A, Adamczyk S, Sun H, Heinonsalo J. 2019. Restriction of plant roots in boreal forest organic soils affects the microbial community but does not change the dominance from ectomycorrhizal to saprotrophic fungi. FEMS Microbiol Ecol 95. https://academic.oup.com/femsec/article/95/9/fiz133/5554003. Last accessed 21/10/2019

Smith GR, Wan J. 2019. Resource-ratio theory predicts mycorrhizal control of litter decomposition. New Phytologist 223:1595–606.

Smith SE, Read DJ. 2008. Mycorrhizal Symbiosis. Academic Press

Soudzilovskaia NA, Bodegom PM van, Terrer C, Zelfde M van’t, McCallum I, McCormack ML, Fisher JB, Brundrett MC, Sá NC de, Tedersoo L. 2019. Global mycorrhizal plant distribution linked to terrestrial carbon stocks. Nat Commun 10:1–10.

Soudzilovskaia NA, van der Heijden MGA, Cornelissen JHC, Makarov MI, Onipchenko VG, Maslov MN, Akhmetzhanova AA, van Bodegom PM. 2015. Quantitative assessment of the differential impacts of arbuscular and ectomycorrhiza on soil carbon cycling. New Phytol 208:280–93.

Steidinger BS, Crowther TW, Liang J, Nuland MEV, Werner GDA, Reich PB, Nabuurs G, de-Miguel S, Zhou M, Picard N, Herault B, Zhao X, Zhang C, Routh D, Peay KG. 2019. Climatic controls of decomposition drive the global biogeography of forest-tree symbioses. Nature 569:404.

Sterkenburg E, Clemmensen KE, Ekblad A, Finlay RD, Lindahl BD. 2018. Contrasting effects of ectomycorrhizal fungi on early and late stage decomposition in a boreal forest. The ISME Journal 12:2187–97.

Subke J-A, Voke NR, Leronni V, Garnett MH, Ineson P. 2011. Dynamics and pathways of autotrophic and heterotrophic soil CO2 efflux revealed by forest girdling. Journal of Ecology 99:186–93.

Taylor MK, Lankau RA, Wurzburger N. 2016. Mycorrhizal associations of trees have different indirect effects on organic matter decomposition. J Ecol 104:1576–84.

Tedersoo L, Bahram M. 2019. Mycorrhizal types differ in ecophysiology and alter plant nutrition and soil processes. Biological Reviews 94:1857–80.

Teste FP. 2008. Role of mycorrhizal networks in dry Douglas-fir forests. University of British Columbia https://open.library.ubc.ca/cIRcle/collections/ubctheses/24/items/1.0066342. Last accessed 21/10/2020

Teste FP, Karst J, Jones MD, Simard SW, Durall DM. 2006. Methods to control ectomycorrhizal colonization: effectiveness of chemical and physical barriers. Mycorrhiza 17:51–65.

Teste FP, Simard SW, Durall DM, Guy RD, Jones MD, Schoonmaker AL. 2009. Access to mycorrhizal networks and roots of trees: importance for seedling survival and resource transfer. Ecology 90:2808–22.

Tisserant E, Malbreil M, Kuo A, Kohler A, Symeonidi A, Balestrini R, Charron P, Duensing N, Frey NF dit, Gianinazzi-Pearson V, Gilbert LB, Handa Y, Herr JR, Hijri M, Koul R, Kawaguchi M, Krajinski F, Lammers PJ, Masclaux FG, Murat C, Morin E, Ndikumana S, Pagni M, Petitpierre D, Requena N, Rosikiewicz P, Riley R, Saito K, Clemente HS, Shapiro H, Tuinen D van, Bécard G, Bonfante P, Paszkowski U, Shachar-Hill YY, Tuskan GA, Young JPW, Sanders IR, Henrissat B, Rensing SA, Grigoriev IV, Corradi N, Roux C, Martin F. 2013. Genome of an arbuscular mycorrhizal fungus provides insight into the oldest plant symbiosis. PNAS 110:20117–22.

Veen GF (Ciska), Freschet GT, Ordonez A, Wardle DA. 2015. Litter quality and environmental controls of home-field advantage effects on litter decomposition. Oikos 124:187–95.

Verbruggen E, Pena R, Fernandez CW, Soong JL. 2017. Chapter 24 - Mycorrhizal interactions with saprotrophs and impact on soil carbon storage. In: Mycorrhizal Mediation of Soil. Elsevier. pp 441–60. https://www.sciencedirect.com/science/article/pii/B9780128043127000243

van der Wal A, Geydan TD, Kuyper TW, de Boer W. 2013. A thready affair: linking fungal diversity and community dynamics to terrestrial decomposition processes. FEMS Microbiol Rev 37:477–94.

Wang Y, Li FY, Song X, Wang X, Suri G, Baoyin T. 2020. Changes in litter decomposition rate of dominant plants in a semi-arid steppe across different land-use types: Soil moisture, not home-field advantage, plays a dominant role. Agriculture, Ecosystems & Environment 303:107119.

Wickham H. 2016. ggplot2: Elegant graphics for data analysis. Springer-Verlag New York https://CRAN.R-project.org/package=ggplot2

Wickham H, Francois R, Henry L, Müller K. 2017. dplyr: A grammar of data manipulation. https://CRAN.R-project.org/package=dplyr https://CRAN.R-project.org/package=dplyr

Wiesmeier M, Urbanski L, Hobley E, Lang B, von Lützow M, Marin-Spiotta E, van Wesemael B, Rabot E, Ließ M, Garcia-Franco N, Wollschläger U, Vogel H-J, Kögel-Knabner I. 2019. Soil organic carbon storage as a key function of soils - A review of drivers and indicators at various scales. Geoderma 333:149–62.

Wurzburger N, Brookshire ENJ. 2017. Experimental evidence that mycorrhizal nitrogen strategies affect soil carbon. Ecology 98:1491–7.

Xu J, Liu S, Song S, Guo H, Tang J, Yong JWH, Ma Y, Chen X. 2018. Arbuscular mycorrhizal fungi influence decomposition and the associated soil microbial community under different soil phosphorus availability. Soil Biology and Biochemistry 120:181–90.

Zak DR, Pellitier PT, Argiroff W, Castillo B, James TY, Nave LE, Averill C, Beidler KV, Bhatnagar J, Blesh J, Classen AT, Craig M, Fernandez CW, Gundersen P, Johansen R, Koide RT, Lilleskov EA, Lindahl BD, Nadelhoffer KJ, Phillips RP, Tunlid A. 2019. Exploring the role of ectomycorrhizal fungi in soil carbon dynamics. New Phytologist 223:33–9.

Zhu W, Ehrenfeld JG. 1996. The effects of mycorrhizal roots on litter decomposition, soil biota, and nutrients in a spodosolic soil. Plant Soil 179:109–18.

