## Supplements for "Ectomycorrhizas accelerate decomposition to a greater extent than arbuscular mycorrhizas in a northern deciduous forest"

Title: Ectomycorrhizas accelerate organic matter decomposition to a greater extent than arbuscular mycorrhizas in a northern deciduous forest

Authors: Alexis Carteron, Fabien Cichonski & Etienne Laliberté

Content: Tables S1 to S3 and Figures S1 to S10.

**Table S1.** For each plot, species basal area (m^2^ ha^-1^) and their mycorrhizal strategy (AM: Arbuscular mycorrhizal, EcM: Ectomycorrhizal). Only individuals of 5 cm diameter and more at breast height were measured.

| Species  Plot | *Acer saccharum*  (AM) | *Fagus grandifolia*  (EcM) | *Acer pensylvanicum*  (AM) | *Acer rubrum*  (AM) | *Betula papyrifera*  (EcM) |
| --- | --- | --- | --- | --- | --- |
| AM plot 1 | 20.3 | 1.73 | 1.12 | 0 | 0 |
| AM plot 2 | 28.3 | 2.6 | 0 | 0 | 0 |
| AM plot 3 | 29.1 | 2.8 | 1.4 | 0 | 0 |
| AM plot 4 | 31.5 | 3.24 | 0.18 | 4 | 0 |
| AM plot 5 | 37.6 | 2.58 | 0 | 0 | 0 |
| EcM plot 1 | 6.89 | 26.2 | 0 | 0 | 4.36 |
| EcM plot 2 | 1.21 | 38.7 | 0.25 | 0.24 | 0 |
| EcM plot 3 | 2.8 | 23.8 | 3.26 | 0 | 0 |
| EcM plot 4 | 14.5 | 23.1 | 0.05 | 0 | 1.44 |
| EcM plot 5 | 3.67 | 29.6 | 0.06 | 0 | 2.94 |

**Table S2.** Effects tests from the full model simultaneously explaining decomposition in arbuscular mycorrhizal and ectomycorrhizal forests.

| Fixed effects | num  DF | den  DF | *F*-value | *P*-value |
| --- | --- | --- | --- | --- |
| (Intercept) | 1 | 206 | 669.3039 | **<0.0001** |
| Soil provenance | 1 | 206 | 5.6962 | **0.0179** |
| Horizon | 2 | 206 | 1537.0702 | **<0.0001** |
| Time | 1 | 206 | 969.2793 | **<0.0001** |
| Mycorrhizal exclusion | 1 | 206 | 24.2292 | **<0.0001** |
| Forest of residence | 1 | 206 | 19.7097 | **<0.0001** |
| Soil provenance × Horizon | 2 | 206 | 4.2256 | **0.0159** |
| Soil provenance × Time | 1 | 206 | 1.3101 | 0.2537 |
| Horizon × Time | 2 | 206 | 11.4403 | **<0.0001** |
| Mycorrhizal exclusion × Forest of residence | 1 | 206 | 0.1923 | 0.6615 |
| Horizon × Mycorrhizal exclusion | 2 | 206 | 4.4747 | **0.0125** |
| Horizon × Forest of residence | 2 | 206 | 2.1804 | 0.1156 |
| Time × Mycorrhizal exclusion | 1 | 206 | 6.5703 | **0.0111** |
| Time × Forest of residence | 1 | 206 | 6.9274 | **0.0091** |
| Soil provenance × Horizon × Time | 2 | 206 | 3.6095 | **0.0288** |
| Horizon × Mycorrhizal exclusion × Forest of residence | 2 | 206 | 0.7767 | 0.4613 |
| Time × Mycorrhizal exclusion × Forest of residence | 1 | 206 | 0.1514 | 0.6976 |
| Horizon × Time × Mycorrhizal exclusion | 2 | 206 | 0.2721 | 0.7620 |
| Horizon × Time × Forest of residence | 2 | 206 | 1.8736 | 0.1562 |
| Horizon × Time × Mycorrhizal exclusion × Forest of residence | 2 | 206 | 0.0139 | 0.9862 |

**Table S3.** Observed values of initial litter chemistry of the upper three horizons (litter - L, fragmented - F, humic - H) of arbuscular mycorrhizal (AM) and ectomycorrhizal (EcM) forest. Means and standard deviations are shown (*n* = 5).

|  | Horizon | AM-dominated  forest | Standard  deviation | EcM-dominated  forest | Standard  deviation |
| --- | --- | --- | --- | --- | --- |
| Total C (%) | L | 46.61 | 0.57 | 47.69 | 0.43 |
|  | F | 44.43 | 1.10 | 46.74 | 0.90 |
|  | H | 41.07 | 2.64 | 46.01 | 1.54 |
| Total N (%) | L | 2.08 | 0.26 | 1.75 | 0.12 |
|  | F | 2.17 | 0.10 | 2.10 | 0.15 |
|  | H | 2.23 | 0.13 | 2.30 | 0.13 |
| Hemicellulose (%) | L | 10.85 | 1.32 | 11.30 | 0.85 |
|  | F | 8.17 | 0.85 | 9.61 | 0.92 |
|  | H | 5.70 | 0.90 | 6.68 | 0.98 |
| Cellulose (%) | L | 16.13 | 1.24 | 19.35 | 0.97 |
|  | F | 11.73 | 1.38 | 14.90 | 1.70 |
|  | H | 10.69 | 1.25 | 11.31 | 1.15 |
| Lignin (%) | L | 19.80 | 1.13 | 22.69 | 1.78 |
|  | F | 23.10 | 2.88 | 26.45 | 3.35 |
|  | H | 19.19 | 2.80 | 24.03 | 2.60 |

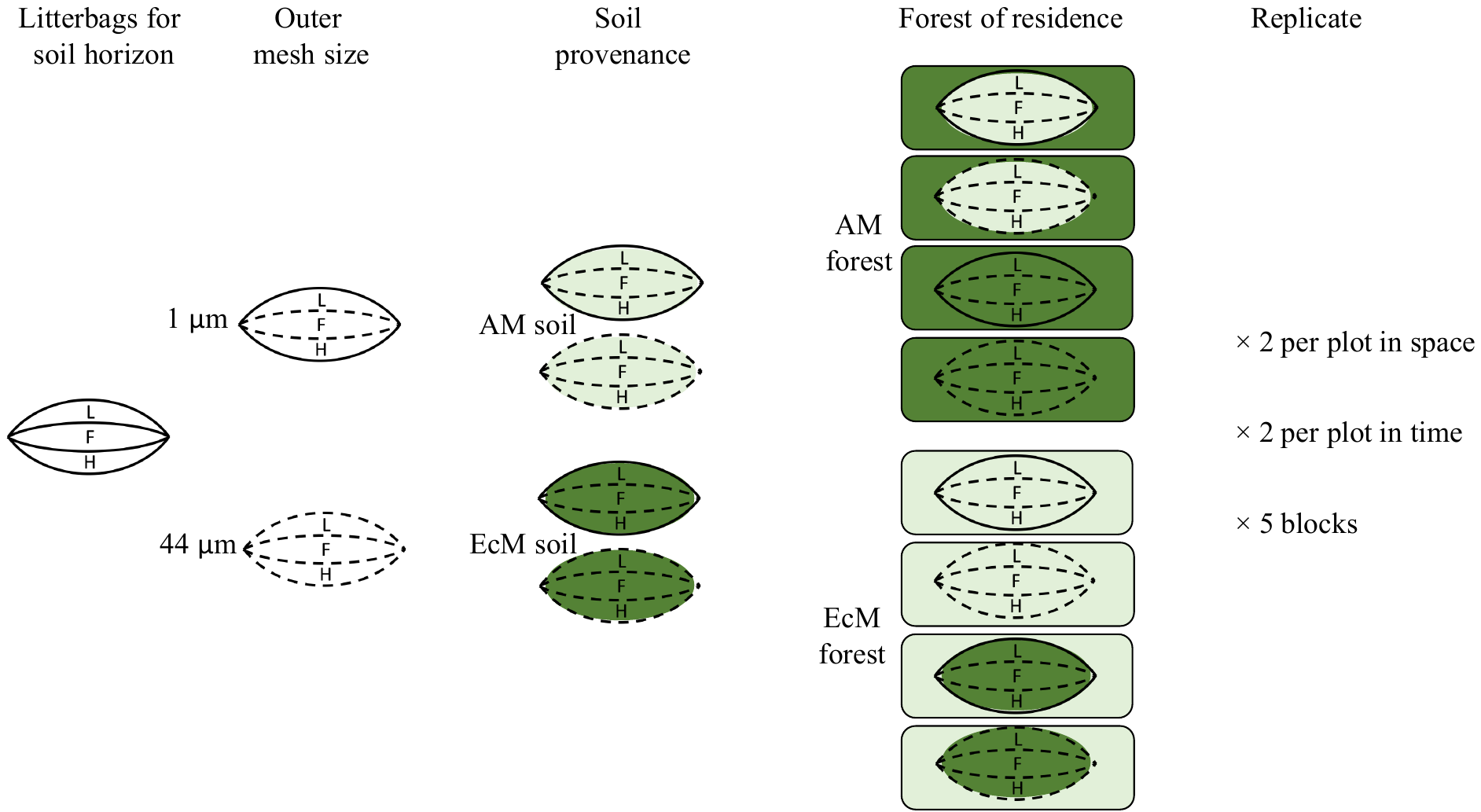

**Figure S1.** Factorial design of the reciprocal transplant experiment. There were 160 litterbags composed of three horizons (L, F, H, standing for litter, fragmented and humic horizon respectively) for a total of 480 incubated soil samples.

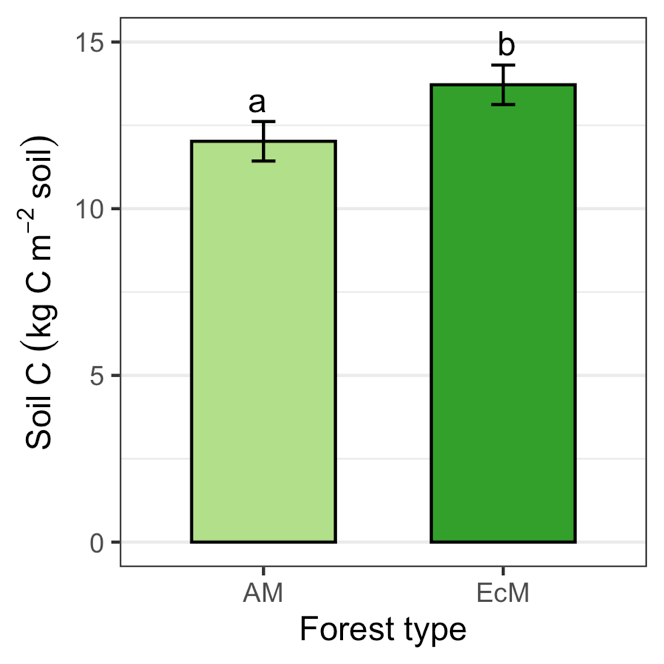

**Figure S2.** Stocks of organic carbon in the upper 20 cm of soil in arbuscular mycorrhizal (AM) and ectomycorrhizal (EcM) forest types. Stocks differed significantly (one-way analysis of variance, *P* < 0.001). Means ± 1 SE are shown (*n* = 5).

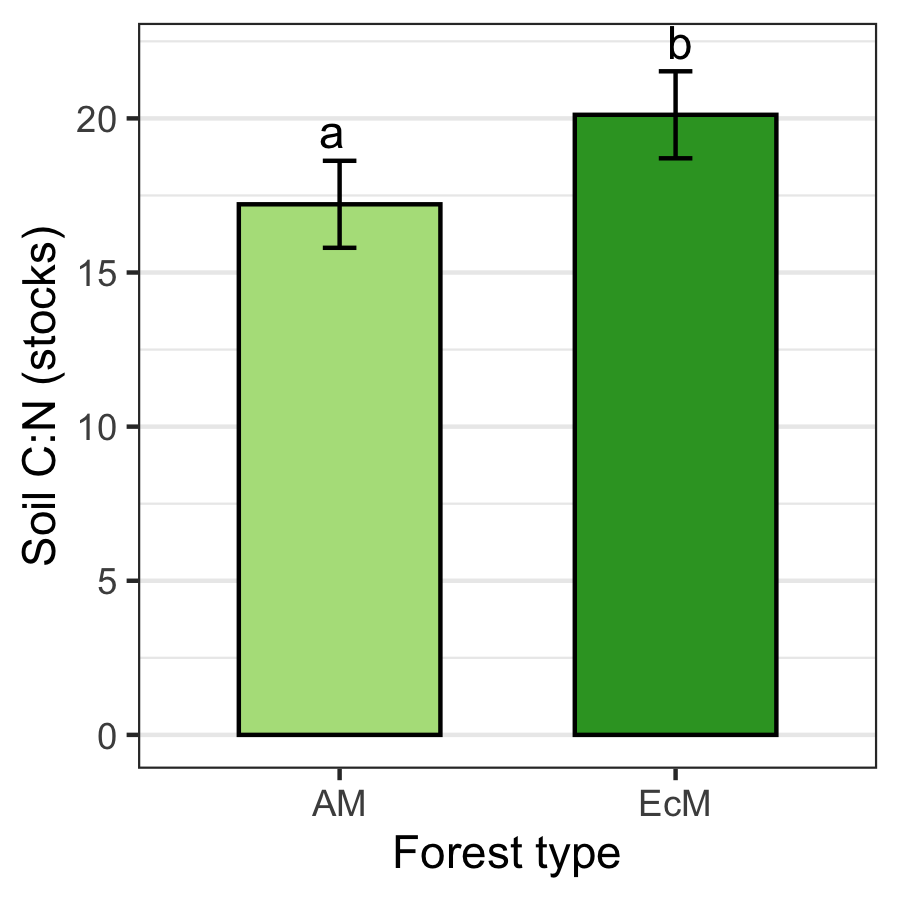

**Figure S3.** C:N ratio in the upper 20 cm differed significantly (one-way analysis of variance, *P* = 0.024) between AM and EcM forests. Means ± 1 SE are shown (*n* = 5).

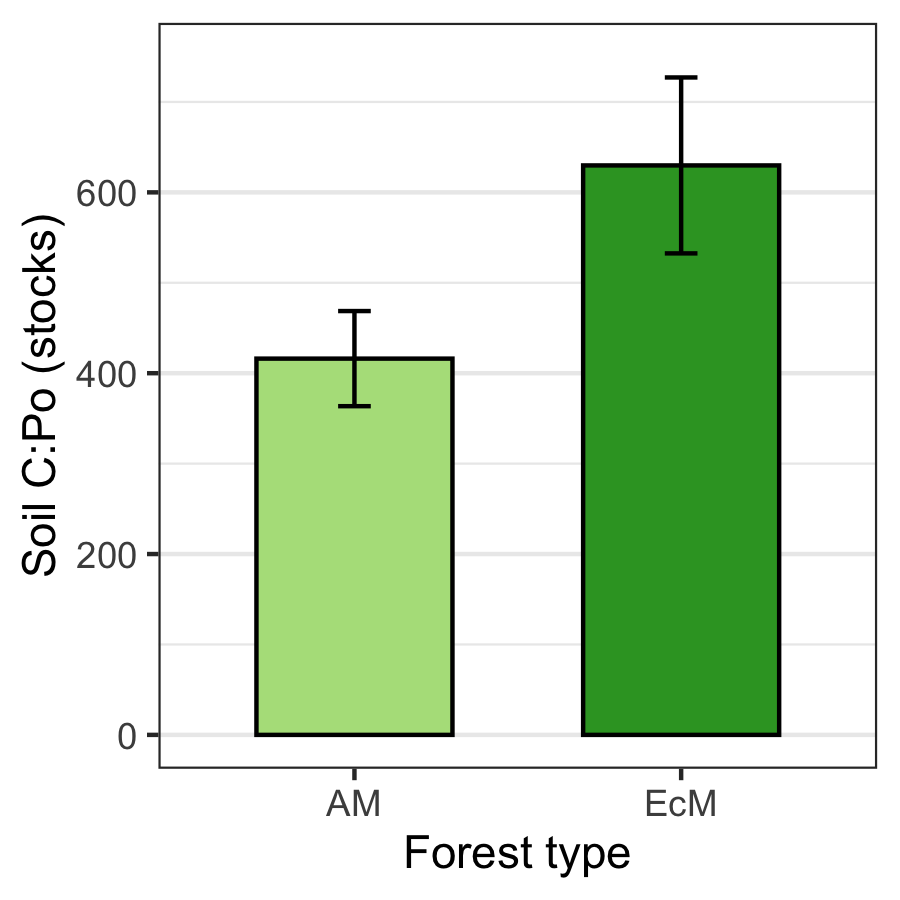

**Figure S4.** C:Po ratio in the upper 20 cm differed but not significantly (one-way analysis of variance, *P* > 0.05) between AM and EcM forests. Means ± 1 SE are shown (*n* = 5).

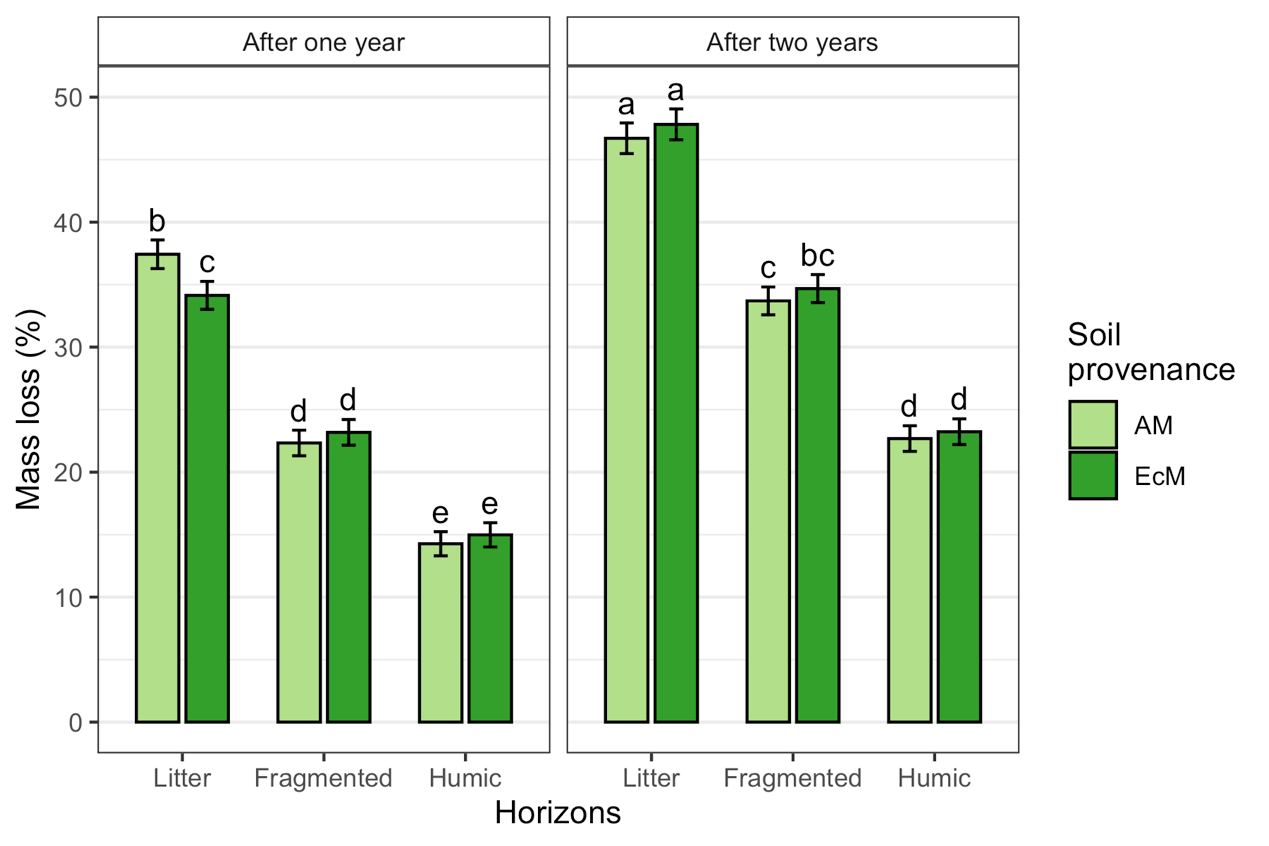
**Figure S5.** Mass loss after one and two years of the three upper horizons originating from arbuscular mycorrhizal (AM) or ectomycorrhizal (EcM) forests. Means ± 1 SE are shown (*n* = 20). Multiple comparison using Tukey's honestly significant difference post-hoc test, different letters indicates significant differences (*P*-value < 0.05).

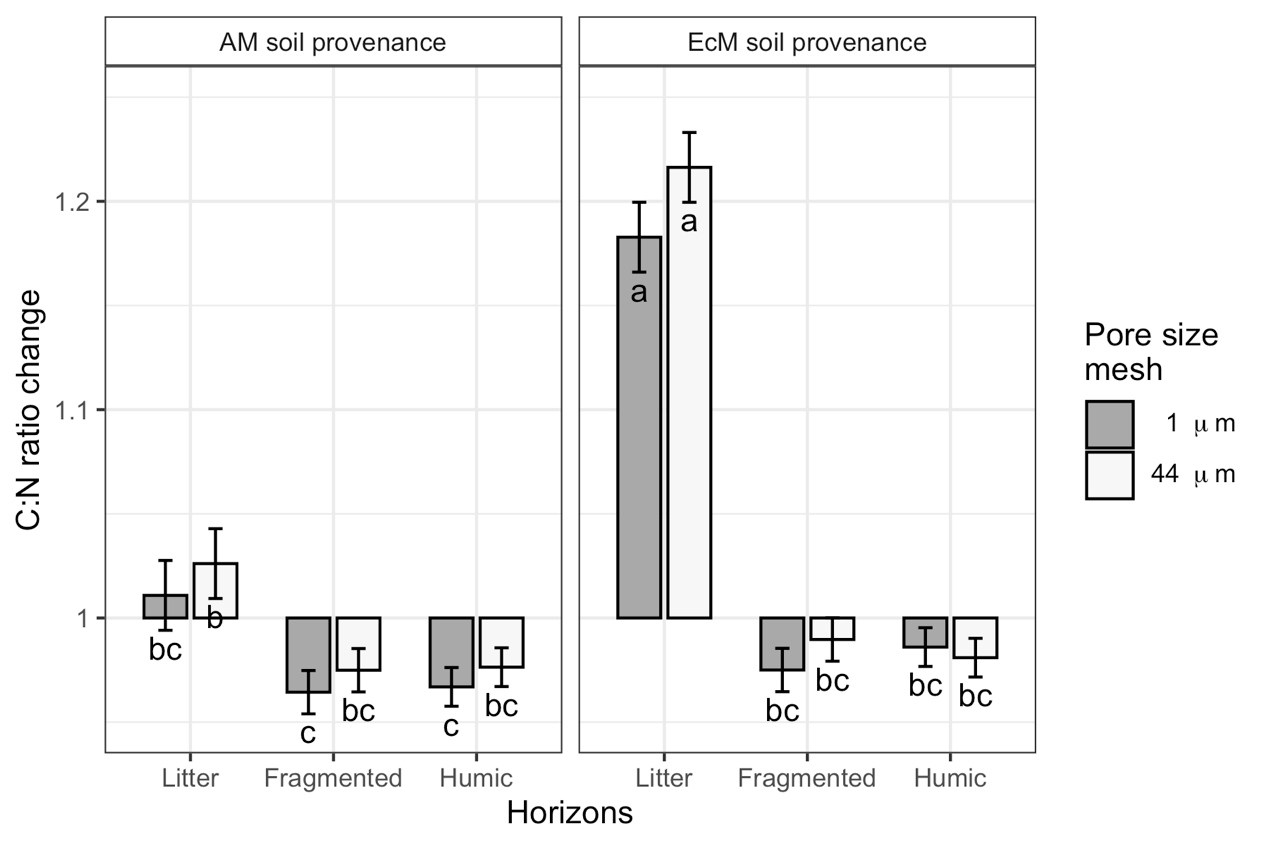

**Figure S6.** Changes in C:N ratio of the three upper horizons originating from the arbuscular mycorrhizal (AM) or ectomycorrhizal (EcM) forests in litterbags with pore size mesh of 1 μm (grey bars) and 44 μm (white bars). Means ± 1 SE are shown (*n* = 20). Multiple comparison using Tukey's honestly significant difference post-hoc test, different letters indicates significant differences (*P*-value < 0.05).

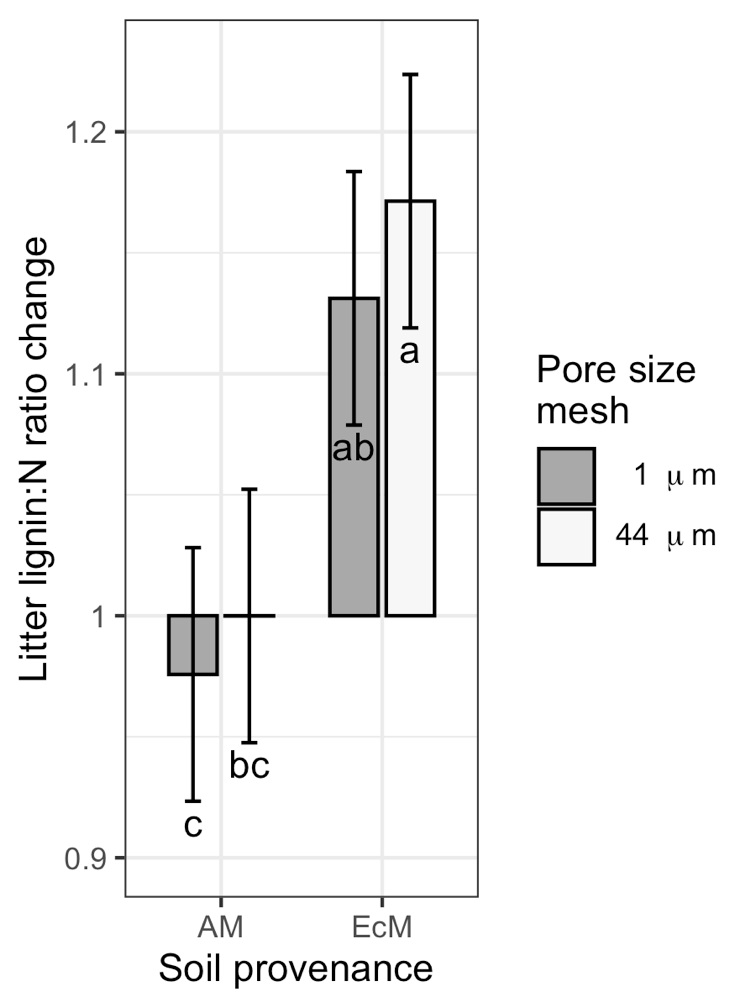

**Figure S7.** Changes in lignin:N ratio in litter (i.e. L horizon) incubated arbuscular mycorrhizal (AM) or ectomycorrhizal (EcM) forests in litterbags with pore size mesh of 1 μm (grey bars) and 44 μm (white bars).. Means ± 1 SE are shown (*n* = 20). Multiple comparison using Tukey's honestly significant difference post-hoc test, different letters indicates significant differences (*P*-value < 0.05).

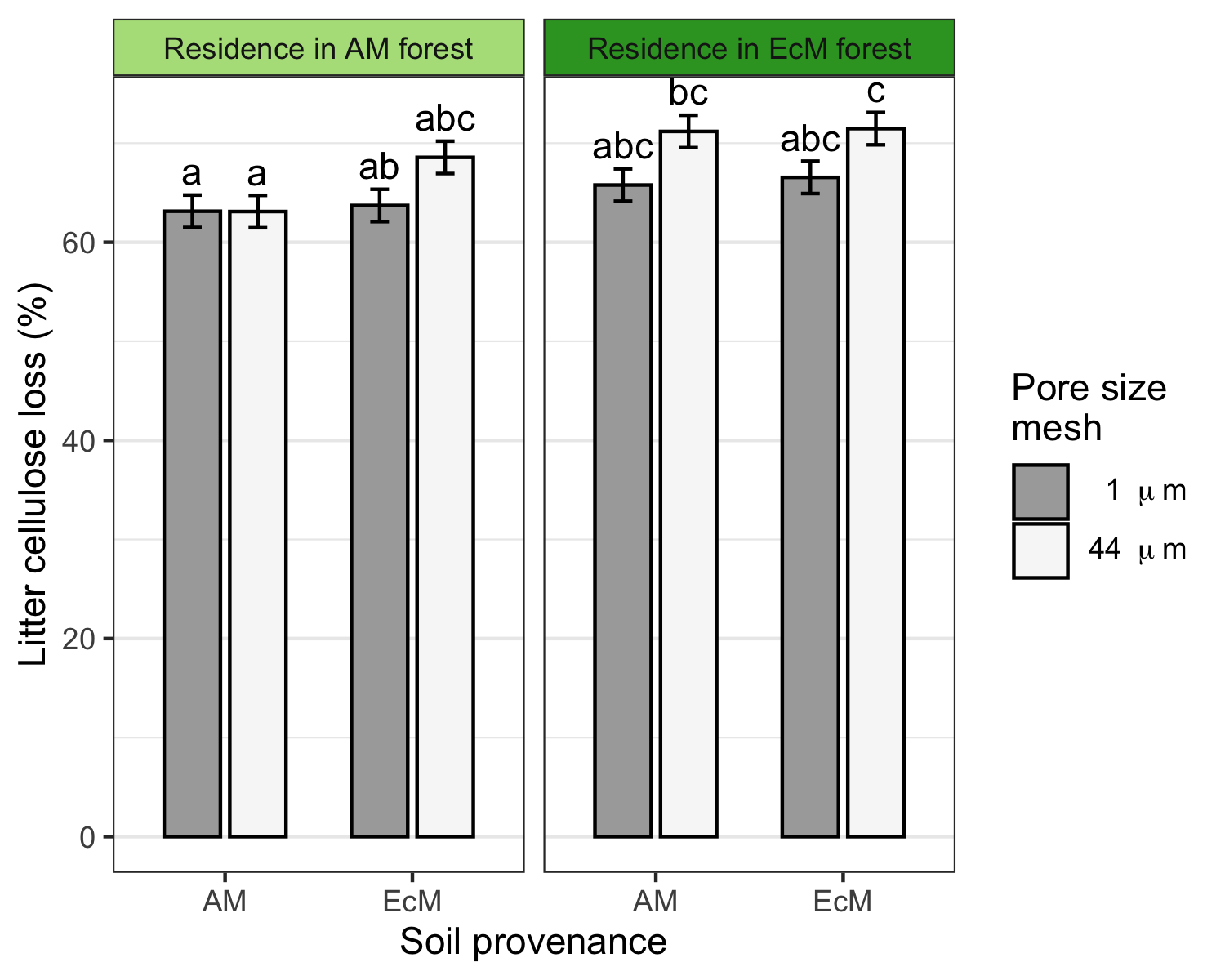

**Figure S8.** Loss of cellulose in litter (i.e. L horizon) incubated for two years in arbuscular mycorrhizal (AM) or ectomycorrhizal (EcM) forests in litterbags with pore size mesh of 1 μm (grey bars) and 44 μm (white bars). Means ± 1 SE are shown (*n* = 20). Multiple comparison using Tukey's honestly significant difference post-hoc test, different letters indicates significant differences (*P*-value < 0.05).

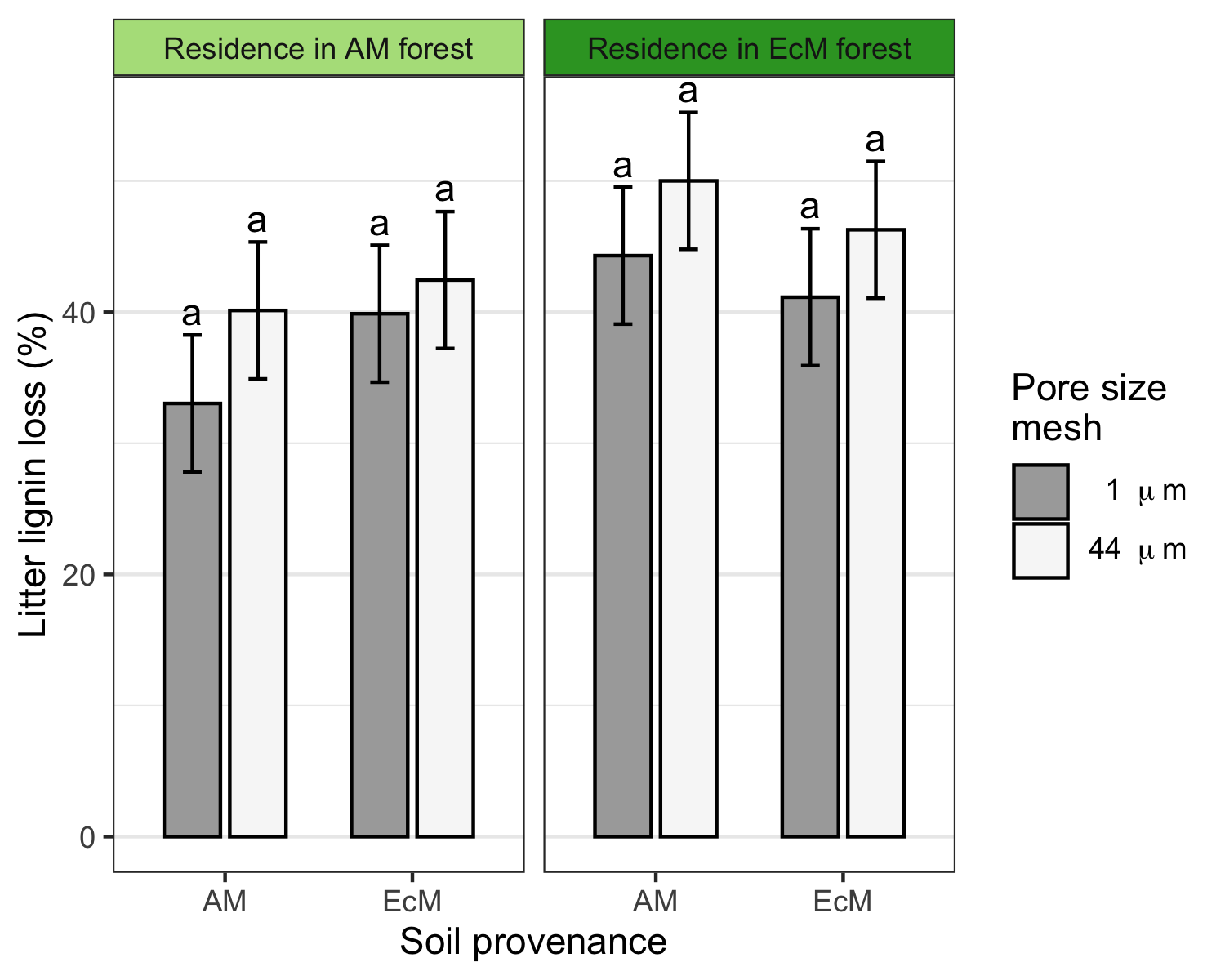

**Figure S9.** Loss of lignin in litter (i.e. L horizon) incubated for two years in arbuscular mycorrhizal (AM) or ectomycorrhizal (EcM) forests in litterbags with pore size mesh of 1 μm (grey bars) and 44 μm (white bars). Means ± 1 SE are shown (*n* = 20). Multiple comparison using Tukey's honestly significant difference post-hoc test, different letters indicates significant differences (*P*-value < 0.05).

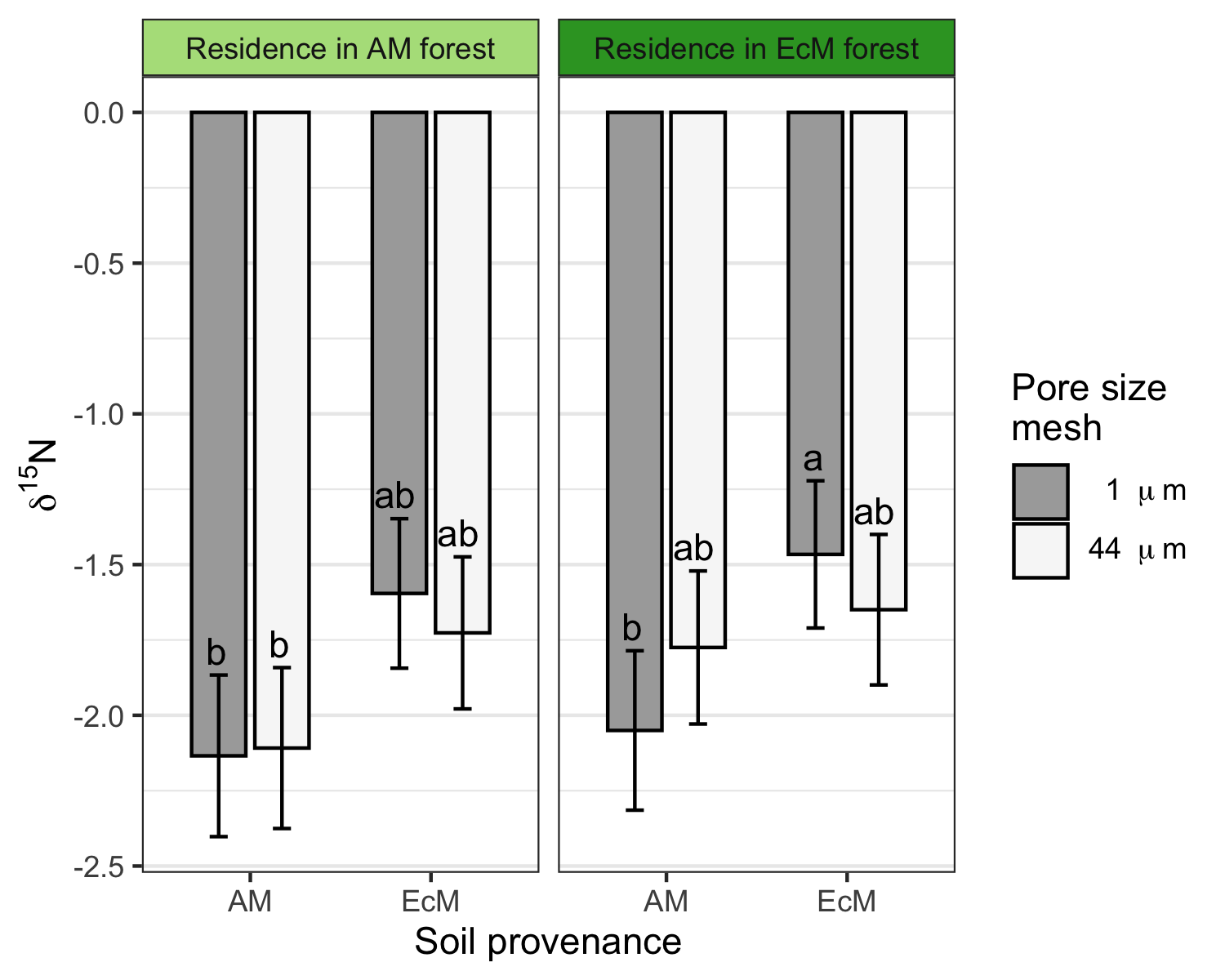
**Figure S10.** Change in ^15^N in fragmented horizons after two years of incubation in arbuscular mycorrhizal (AM) or ectomycorrhizal (EcM) forests in litterbags with pore size mesh of 1 μm (grey bars) and 44 μm (white bars). Means ± 1 SE are shown (*n* = 5). Multiple comparison using Tukey's honestly significant difference post-hoc test, different letters indicates significant differences (*P*-value < 0.05).
